# Tannins and gut health in broilers: Effects of a blend of chestnut and quebracho tannins on gut health and performance of broiler chickens

**DOI:** 10.1101/2021.07.27.454004

**Authors:** Enzo A. Redondo, Leandro M. Redondo, Octavio A. Bruzzone, Juan M. Diaz-Carrasco, Claudio Cabral, Victorino Garces, Maximo Liñeiro, Mariano E. Fernandez-Miyakawa

## Abstract

Consumer demands and increased regulations on the use of antimicrobials in farm animals accentuated the need to develop strategies to replace antimicrobial growth promoters (AGPs) in food-producing animals. The present study evaluates the productive and gut health outcomes during the implementation of AGPs free programs based on the inclusion of a tannin blend compared with AGPs based program under commercial conditions. In the first trial, 6 farms were randomly assigned to AGP or tannin-based programs. In a second trial, both programs were applied simultaneously in one farm and the results were studied over 1 year. Although productive results from both trials were similar among treatments, evaluations of gut health indicators show improvements in the tannins treated flocks. Frequency and severity of intestinal gross lesions were reduced in jejunum (42% vs 23%; p<0.05 – 1.37 vs. 0.73; p<0.01, respectively) and ileum (25% vs. 10%; p<0.0.5 – 1.05 vs. 0.58; p<0.01) in tannins treated birds. Results from 16S studies, show that cecal microbiota diversity was not differentially affected by AGPs or tannins, but changes in the relative abundance of certain taxa were described, including *Lactobacillus* and *Bifidobacterium* groups. Additional evaluations using an *in vivo* model for *C. perfringens* necrotic enteritis showed that tannins treated birds had reduced incidence of gross lesions in jejunum (43.75 vs. 74.19%; p<0.01) and ileum (18.75% vs. 45.16%; p<0.05) compared with control. These results suggest that AGPs can be replaced by tannins feed additives, and contribute in the implementation of antimicrobial-free programs in broilers without affecting health or performance.

## INTRODUCTION

Restrictions on the use of antimicrobials have led to an increase of intestinal health problems in broiler chickens and thus reduced profitability for farmers [1]. A clear example is the increased incidence of *C. perfringens* necrotic enteritis (NE) in countries where the use of antimicrobial growth promoters (AGPs) has been banned [2, 3]. This situation not only highlights an excessive dependency of modern animal production on antimicrobials, but also prompted the search for cost and biologically effective alternatives [4, 5]. A large and diverse group of potential alternatives has been investigated and developed over the last years to replace AGPs. Considering that the ideal AGP alternative should improve productive efficiency and promote animal health, plant phytochemicals represent a group of promising candidates [5]. The described potential beneficial effects of phytochemicals include maintenance of gut integrity, promotion of beneficial bacteria growth and reduction of negative consequences of bacterial infections [5–7]. Phytochemicals are natural bioactive compounds derived from plant tissues, which include a wide variety of molecules such as tannins, terpenoids, alkaloids and flavonoids, many of which have been found to have an extensive arrangement of biological activities including antimicrobial, antioxidant and anti-inflammatory properties [4–6].

Among phytochemicals, tannins stand out for their role as alternative to AGPs in poultry production [7]. In broiler chickens, several tannins have proven to improve growth performance [8] and reduce the detrimental effects of *C. perfringens* infection [9]. Previous works from our group, describe that hydrolysable tannins derived from extracts of chestnut tree (*Castanea sativa*) and condensed tannins obtained from quebracho trees (*Schinopsis lorenzii*) have antimicrobial and antitoxin activities against *C. perfringens* [10–11]. In fact, it has been shown that they improve gut health and productive parameters under experimental [12–13] and commercial conditions [14]. In addition, available data suggest that their use does not contribute to selection and spread of antimicrobial resistance, since pathogenic bacteria such as *C. perfringens* do not generate resistance against these tannins even after long term exposure [11].

To overcome limitations to study the application of potential AGP alternatives, multiple approaches are needed to identify products with high effectiveness rate to reduce antimicrobial needs in poultry industry. Besides experimental trials describing the effects of AGP alternatives and their potential mode of action, efficacy should be contrasted in field/commercial trials under different challenge conditions. However, only a few studies describing the impact of AGP-free programs in commercial poultry production conditions are available in the scientific literature, and these works reported variable results [15–18]. The objective of the present report was to study the consequences of the implementation of AGP free programs based on the inclusion of a blend of chestnut and quebracho tannins, and how this additive can affect broiler productive performance, intestinal health and cecal microbiota in comparison with conventional AGP based programs. In order to fulfil this goal, three studies with different approaches were conducted: (1) a trial comparing the effects on different commercial farms at the same time; (2) the follow-up of one farm during an entire productive year; and (3) an experimental trial using a *C. perfringens* challenge model in broiler chickens.

## MATERIALS AND METHODS

### Experiment 1 (Study design)

A prospective study was conducted in 6 commercial broiler chicken farms located in the province of Buenos Aires, Argentina. Farms were selected within a radius of 200 km from the Instituto de Patobiologia Veterinaria, CICVyA – INTA among voluntary producers. To be included in the study, each farm was required to have 2-6 houses, with similar stocking densities, surface areas, feeding systems, water equipment and ventilation systems. The capacity of the houses ranged from 10,000 to 20,000 broiler chickens per flock. Selected farms were working under contract with the same integrated chicken company, sharing same hatchery, feed mill and slaughterhouse as well as management practices. During this study, chickens were fed in four phases: a pre-starter diet (days 1 – 14), starter diet (days 15 – 21), finisher diet (day 22 – 35), and withdrawal diet (day 36 – clear), nutritional information is summarized in Table 1. Each farm was randomly assigned to one of two treatments: (1) conventional AGP program, or (2) tannins-based AGP free program. Birds raised under conventional protocol of the company were fed with commercial diets supplemented with Enramycin (10 g/Tn) as AGP, while in the tannins protocol broiler feed was supplemented with a blend of chestnut and quebracho tannins (SILVAFEED/NUTRI-P 1 Kg/Ton). Both groups were raised similarly, following the conventional management practices as per their feed mill and hatchery guidelines, basically Cobb management handbooks (CoBB-Vantress, 2013). No specific recommendations pertaining to water acidification and brooding were given, and therapeutic antimicrobials or anticoccidial drugs were prescribed by the company veterinarian when needed. Each farm was visited at 1, 21 and 42 days bird’s age. Visits on day 1 were organized to control chicken’s general health conditions at the beginning of every producing period. Parameters such as quality of anticoccidial vaccination, based on the percentage of chicks with the presence of pink dye on the head, crop filling and cloacal temperature were monitored. Results from the 24 h-visits will not be presented in this paper.

**Table 1:** Dietary composition and nutrient levels

| **Ingredients (%)** | **Pre-starter** | **Starter** | **Grower** | **Withdrawal feed** |
| --- | --- | --- | --- | --- |
| Corn | 63,124 | 58,573 | 58,752 | 61,661 |
| Soybean meal | 25,658 | 25,784 | 23,521 | 21,550 |
| Corn gluten meal | 3,774 | - | - | - |
| Meat meal | 3,500 | 4,873 | 3,500 | 3,500 |
| Soy oil | 1,000 | 1,775 | 2,100 | 2,200 |
| Mono/bicalcium phosphate | 0,624 | - | 0,265 | 0,115 |
| Lysine 50% | 0,582 | 0,303 | 0,205 | 0,195 |
| Salt | 0,402 | 0,231 | 0,255 | 0,254 |
| Liquid methionine | 0,200 | - | - | 0,200 |
| Shell poder | 0,182 | - | 0,214 | 0,174 |
| Powder methionine | 0,149 | 0,313 | 0,229 | 0,043 |
| Choline 75% | 0,100 | 0,090 | 0,080 | 0,070 |
| Mineral premix | 0,100 | 0,100 | 0,080 | 0,040 |
| Vitamin premix | 0,100 | 0,100 | 0,080 | 0,040 |
| Mycotoxin sequestrant | 0,100 | - | - | - |
| Sodium bicarbonate | 0,100 | 0,100 | 0,100 | 0,100 |
| Threonine | 0,095 | 0,078 | 0,039 | 0,005 |
| Anticoccidial | 0,060 | 0,060 | 0,060 | 0,060 |
| Protease | 0,020 | - | - | - |
| Phytase | 0,020 | 0,020 | 0,020 | 0,020 |
| Antioxidant | 0,010 | - | - | 0,010 |
| Extruded soybean | - | 4,500 | 5,500 | 5,500 |
| Wheat | - | 3,000 | 4,900 | 4,263 |
| **Calculated nutrients** | **Pre-starter** | **Starter** | **Grower** | **Withdrawal feed** |
| Metabolizable energy (Kcal/kg) | 3.015 | 3.110 | 3.160 | 3.200 |
| Crude protein (%) | 21,100 | 20,370 | 19,250 | 19,080 |
| Dig. Lysine (%) | 1,200 | 1,100 | 0,990 | 0,970 |
| Dig. Methionine (%) | 0,600 | 0,578 | 0,481 | 0,451 |
| Dig. Met + Cys (%) | 0,900 | 0,836 | 0,733 | 0,717 |
| Total phosphorus (%) | 0,750 | 0,720 | 0,684 | 0,646 |
| Available phosphorus (%) | 0,480 | 0,440 | 0,420 | 0,390 |
| Calcium (%) | 0,850 | 0,840 | 0,800 | 0,750 |

### Experiment 2 (Study design)

Based on productive performance, structural and sanitary conditions, one of the farms included in “Experiment 1” was selected to compare the global outcome of using AGPs or tannins based-programs on broiler performance and health over a one-year period. In the selected farm, each of six tunnel ventilated broiler houses (∼20,000 birds) was randomly assigned to one of two treatments: (1) conventional AGP program (AGP rotation: Bacitracin 50 g/Tn, Avilamycin 10 g/Tn, Enramycin 10 g/Tn), or, (2) tannins-based AGP free-program (SILVAFEED/NUTRI-P 1 Kg/Ton). This assignment was kept during 6 production cycles over a 5- or 7-week period according to commercial requirements. A total of 720,000 mixed sex birds were included in this trial. General management practices other than antimicrobial use were like those described for “Experiment 1”. Similarly, during each productive cycle this farm was visited at 1, 21 and 35/42 days. All barns were monitored, and general health of the birds was controlled as described for experiment 1.

### Data and sample collection from field trials

#### Productive performance

In both experiments, zootechnical performances of the flocks included in this study were provided by the poultry producer. On each house included in field trials, 50 birds were randomly selected to determine weekly body weight (BW), additional body weight records were obtained from slaughterhouse. Mortality was obtained weekly and feed consumption was obtained at the conclusion of each cycle. Feed conversion (FCR) was calculated as the feed to gain ratio. The gain, feed intake, and feed conversion were corrected for dead birds. European Poultry Efficiency Factor (EPEF) values were calculated for overall growth period using the following standard formula: EPEF = [(BW in Kg x % Livability)/ (Age in days x FCR)] x 100.

#### Necropsy and sample collection

A similar methodology was used for both experiments. On day 21, 10 healthy male birds per barn were randomly selected. Birds were euthanized by cervical dislocation and complete necropsy was performed immediately for examination of gross lesions. During necropsies, the small intestinal segments and ceca were carefully observed, and gross lesions were recorded and scored blindly by two experienced pathologists as described by Cooper and Songer [19]. Footpad lesions were determined by visual inspection and palpation and classified using a 5-point score system based on the presence, size and severity of lesions [20]. Segments of 2-cm in length from the mid-points of the duodenum, jejunum, and ileum were cut, flushed with cold saline, and immediately preserved in 10% phosphate-buffered formalin for histomorphological analysis [21]. Additionally, during experiment 2, cecal contents were transported at 4°C and stored at -80°C for DNA extraction and determination of cecal microbiota. During rearing periods, all groups were routinely supervised, and clinical examinations were performed regularly. Additional visits were conducted if an increased mortality rate was observed by the producer. In those circumstances, necropsies were performed on moribund and dead birds. Finally, birds were visited on the day before slaughter (day 35 or 42). Necropsies, lesion scoring, and intestinal sampling were repeated exactly as described for day 21.

#### Experiment 3

Based on necropsy findings from experiments 1 and 2, further trials under experimental conditions were performed using an *in vivo* infection model for *C. perfringens* necrotic enteritis (NE) in broilers [19].

#### Birds and housing

Experimental NE challenges were performed with 175 male Cobb broiler chickens housed in biosafety level 2 facilities located in the Veterinary and Agriculture Research Center (CICVyA-INTA), with controlled temperature and humidity and automated ventilation system. Birds were obtained as 1-day-old chicks from the same commercial hatchery which provides birds for experiments 1 and 2. On arrival day, birds were randomly divided in 15 groups of 11-12 chicks each. All treatment groups were housed in the same room, placed in pens (1.5 × 1.5 × 0.8 m) made of 0.55 mm wire mesh and hardboard pieces covering the lower part of the mesh to separate them. Commercial wood shavings were used as bedding material and maintained until the end of the trial. Commercial starter feed described before (Table 1) was placed in galvanized steel trays and given ad-libitum, the same for water. For experimental treatments, the commercial feed was mixed with different tannin-based additives (final concentration 1 Kg/Ton), and each group of birds was randomly assigned to one of 5 experimental treatments: (1) negative control: feed without tannin additives, NE unchallenged; (2) positive control: feed without tannins additives, NE challenged; (3) chestnut: feed with chestnut based additives, NE challenged; (4) quebracho: feed with quebracho based additives, NE challenged; (5) mix: commercial mix of chestnut and quebracho (SILVAFEED – NUTRI P) additive, NE challenged).

#### *In vivo* NE model

A *C. perfringens* strain (*cpa+, cpb2-, netB+, TpeL-*) isolated from a natural outbreak of broiler NE was used as inoculum to challenge broilers. The strain was prepared by streaking glycerol aliquots onto blood agar plates and grow under anaerobic conditions (5% H_2_:5% CO_2_:90% N_2_) for 18 h at 37 °C. Then, 1-2 colonies were transferred into 10 ml cooked meat medium (CMM) and further incubated for 18 h at 37 °C. This culture was inoculated in 100 ml fluid thioglycollate broth (FTG) and cultured as before. After 18 hours, the FTG culture was diluted 1:10 in CMM and incubated as before. One hundred ml of the last CMM culture was used as inoculum for 1 L of FTG medium, after 18 h of incubation this culture reached a concentration of 7–9 × 10^8^ cfu/ml. This last FTG culture was used as inoculum and procedure was repeated for each dose used during the challenge (total challenge doses, n=6). On day 15, birds were fasted for 12 h prior to the initial challenge on day 16. Challenge was performed orally by mixing *C. perfringens* FTG culture with commercial feed (1:1 v/w) and given to birds twice a day on days 16, 17 and 18. Uneaten feed was replaced before each subsequent challenge. All groups were routinely supervised and clinical examinations were performed regularly. On day 19, some birds from each group (n=5) were euthanized by cervical dislocation and necropsy was performed immediately for examination of intestinal gross lesions as described for the field trials. To confirm the identity of intestinal lesions, tissue samples were taken from each bird with gross lesions. Samples were kept refrigerated or in buffered formalin solution for both bacteriological and histopathological diagnosis.

#### Microscopic Analysis of Digestive Tracts

Intestinal segments were fixed for at least 48 h in 10% phosphate-buffered formalin for histological analysis. The processing consisted of serial dehydration, clearing, and impregnation with wax. Tissue sections, 5 μm thick (3 cross-sections from each sample), were cut by a microtome and were fixed on slides. A routine staining procedure was carried out using hematoxylin and eosin. Histological analysis of the intestinal samples was done using standard light microscopy (Microscope NIKON ECLIPSE 80i Co., Ltd, Japan), a camera (DS-Fi1c) and a computer-based image analysis system (ImageJ, National Institutes of Health). Villi fusion, presence of coccidia, necrotic debris, capillary congestion, etc. were considered. For histomorphometric studies, well-oriented crypt-villus units were selected in triplicate for each intestinal cross-section for each sample. The criterion for villus selection was based on the presence of intact lamina propria. Villus height (VH, from the tip of the villus to the top of the lamina propria), crypt depth (CD, from the base to the region of transition between the crypt and villus) and the thickness of the muscularis mucosae in the duodenum, jejunum and ileum were determined. Measurements of 10 complete villi for VH and associated crypts for CD were taken from each segment and the average of these values was used for statistical analysis [21].

#### Caecal Microbiota Composition

Total DNA was isolated from 300 mg of cecal contents using QIAamp DNA Stool Mini Kit (Qiagen, Hilden, Germany) following manufacturer instructions. DNA concentration and quality were assessed in NanoDrop ND-1000 spectrophotometer (NanoDrop Technologies, DE, USA). DNA was stored at −20∘ C until further analysis. Amplification of 16S rRNA Gene (V3-V4 region) and high-throughput sequencing was performed as previously described [13]. Microbial composition and diversity were analysed using Quantitative Insights into Microbial Ecology 2 (QIIME2) software [22].

#### Institutional Animal Care and Use Committee (IACUC) Approval

Experimental and field studies presented here were performed in accordance with the ARRIVE guidelines (https://www.nc3rs.org.uk/arrive-guidelines). The experiments were approved by the Institutional Animal Care and Use Committee of the Veterinary and Agriculture Research Center -INTA, under protocol number 20/2010.

#### Estimation of Parameters to Describe Growth

To study the growth, a Gompertz growth curve [23] was fitted to the data and the parameters of the curve (initial weight, growth rate, and final weight) were compared between treatments. Over that base model, we added three more parameters, described in Table S1 (autocorrelation coefficient, error intercept, and error slope) to compensate for measurement errors, and autocorrelation, as described below.

An autoregressive model was used to compensate for the autocorrelation caused by the repeated measures on the same individual, so the growth curve was (equation 1):

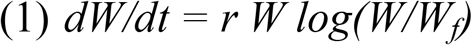

with initial conditions *W_(t=0)_ = W_o_*, being W the weight in grams of the animal, and *r* its growth rate in 1/days, W_o_ is the weight at the beginning of the experiment. Given that the animals were measured repeated times, *W_i_* represents the estimated weight at the measurement time *i*, which was corrected using an autoregressive model as follows (equation 2):

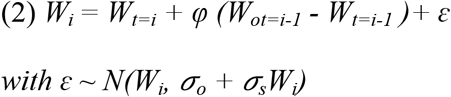

with t being time, i-1 being the time of previous measurement, W_o_ the observed weight, and φ an autoregressive coefficient, ɛ are the errors which are normally distributed with mean zero, as we observed that errors increased with the weight of the animal, then we used a standard deviation which increases linearly with the expected weight, being *σ_o_* the intercept and *σ_s_* its slope.

All the parameters and information indexes of the proposed models were calculated using Markov Chain Monte Carlo (MCMC) methods with the Metropolis-Hastings algorithms. The Monte Carlo calculations were performed with the python programming language and the pymc version 2.4 for Monte Carlo methods [24].

### Statistical Analysis

For field trials the statistical unit was the flock in all analyses, and AGPs treated groups were considered as control for analysis. Performance data were corrected with mortality and subjected to one-way ANOVA analysis. Necrotic enteritis lesion scores were statistically analysed using a two-tailed Fisher exact test to assess the difference between proportions of pathological lesion development between treatment groups in both field trials and the experimental challenge trial. Paired t tests were used for analysis of lesion scores and histomorphometric determinations. Differences were considered significant at P ≤ 0.05.

The relative abundances of bacterial populations were analysed using Statistical Analysis of Metagenomic Profiles (STAMP) software [25]. Relative abundances were compared by White’s test. Correlations between productive efficiency indicators and taxonomic relative abundance at the phylum and genus levels were determined using Spearman correlation coefficients, only r index with p-values<0.05 were considered.

## RESULTS

### Experiment 1

#### Productive performance

No significant differences were observed in live weight (2.621 kg vs. 2.523 kg), feed conversion (1.71 vs. 1.73), mortality (3.9% vs. 3.4%) or EPEF (300 vs. 288) between farms under AGP and tannins-based programs respectively.

#### Necropsy findings and intestinal structure

Frequency and severity of intestinal gross lesions were reduced in birds from tannins treated farms (Table 2). Jejunal gross lesions were present in 23.3% of the necropsied birds compared with the 42.5% in the AGP farms (p<0.05), similar results were observed in the ileum (10% vs 25%, p<0.05), although differences were not significant, duodenal lesions were also reduced (24.4% vs. 30%). Those differences were more evident during the starter period (day 21). Similar results were obtained regarding lesion severity, gross lesion score were reduced in jejunum (0.73 vs 1.37, p<0.01) and ileum (0.58 vs. 1.05, p<0.005), duodenal scores were also reduced although differences were not significant (0.87 vs 1.15). No differences were observed in the frequency of footpad lesions and presence of undigested feed in distal portions of digestive tract. Histomorphological measurements for duodenal, jejunal and ileal sections were similar in both treatments; results from these determinations are presented in Table 3.

**Table 2:** Frequency ans score of intestinal gross lesions^1^ (Experiment 1)

|  | AGPs |  | Tannins |  |
| --- | --- | --- | --- | --- |
|  | Frequency (%) | Score (mean) | Frequency (%) | Score (mean) |
| **Day 21** |  |  |  |  |
| Duodenum | 30 | 1.30 | 28 | 1.08 |
| Jejunum | 35 | 1.35 | 18 * | 0.56 * |
| Ileum | 35 | 1.20 | 12 * | 0.58 * |
| **Day 42** |  |  |  |  |
| Duodenum | 30 | 1.00 | 20 * | 0.62 |
| Jejunum | 50 | 1.40 | 30 | 0.95 |
| Ileum | 15 | 0.90 | 7.50 | 0.60 * |
| **Total** |  |  |  |  |
| Duodenum | 30 | 1.15 | 24.4 | 0.87 |
| Jejunum | 42.5 | 1.37 | 23.3 * | 0.73 ** |
| Ileum | 25 | 1.05 | 10 * | 0.58 ** |

\*: p<0.05; \*\*: p<0.01
<sup>1</sup> Intestinal gross lesion score: 0: no apparent gross lesions; 1: removable fibrin deposit; 2: isolated focal necrosis or ulceration (1 to 5 foci); 3: multiple focal necrosis or ulceration (6 or more foci); 4: extensive areas of necrosis; 5: diffuse necrosis, presence of attached pseudomembrane).

**Table 3:**
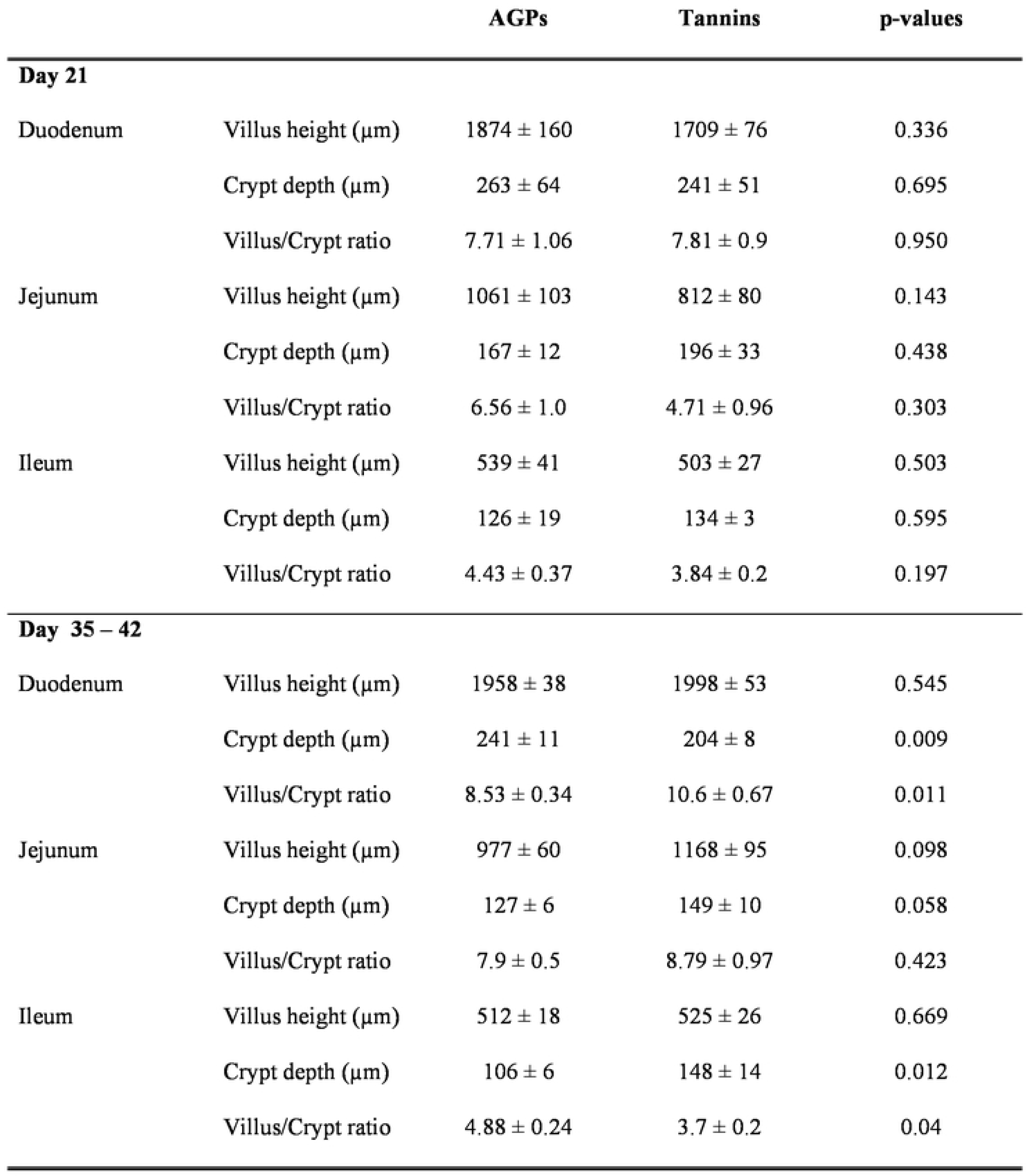
Intestinal histomorphometry (Experiment 1)

|  |  | **AGPs** | **Tannins** | **p-values** |
| --- | --- | --- | --- | --- |
| **Day 21** |  |  |  |  |
| Duodenum | Villus height (µm) | 1874 ± 160 | 1709 ± 76 | 0.336 |
|  | Crypt depth (µm) | 263 ± 64 | 241 ± 51 | 0.695 |
|  | Villus/Crypt ratio | 7.71 ± 1.06 | 7.81 ± 0.9 | 0.950 |
| Jejunum | Villus height (µm) | 1061 ± 103 | 812 ± 80 | 0.143 |
|  | Crypt depth (µm) | 167 ± 12 | 196 ± 33 | 0.438 |
|  | Villus/Crypt ratio | 6.56 ± 1.0 | 4.71 ± 0.96 | 0.303 |
| Ileum | Villus height (µm) | 539 ± 41 | 503 ± 27 | 0.503 |
|  | Crypt depth (µm) | 126 ± 19 | 134 ± 3 | 0.595 |
|  | Villus/Crypt ratio | 4.43 ± 0.37 | 3.84 ± 0.2 | 0.197 |
| **Day 35 – 42** |  |  |  |  |
| Duodenum | Villus height (µm) | 1958 ± 38 | 1998 ± 53 | 0.545 |
|  | Crypt depth (µm) | 241 ± 11 | 204 ± 8 | 0.009 |
|  | Villus/Crypt ratio | 8.53 ± 0.34 | 10.6 ± 0.67 | 0.011 |
| Jejunum | Villus height (µm) | 977 ± 60 | 1168 ± 95 | 0.098 |
|  | Crypt depth (µm) | 127 ± 6 | 149 ± 10 | 0.058 |
|  | Villus/Crypt ratio | 7.9 ± 0.5 | 8.79 ± 0.97 | 0.423 |
| Ileum | Villus height (µm) | 512 ± 18 | 525 ± 26 | 0.669 |
|  | Crypt depth (µm) | 106 ± 6 | 148 ± 14 | 0.012 |
|  | Villus/Crypt ratio | 4.88 ± 0.24 | 3.7 ± 0.2 | 0.04 |

### Experiment 2

#### Productive Performance

Productive outcomes were similar among flocks included in both treatments. No statistically significant differences were observed between AGP and tannin treatments in weekly (Table 4) or final bodyweight either at 5 (1.452 Kg vs. 1.491 Kg, respectively) or 7 weeks (2.538 Kg vs. 2.608 Kg), flock uniformity expressed as CV (12,9% vs. 12,5%), feed conversion (1.767 vs. 1.762) or EPEF values (268 vs. 271). During this experiment and depending on commercial requirements, birds were slaughtered at different ages. Therefore, potential growth and weight gains were estimated by Gompertz equation. According to this model, tannins treated birds will have higher growth potential and growth rate may decrease at a slower rate compared with the AGPs treated groups (Figure 1) (Table S1).

**Figure 1:**
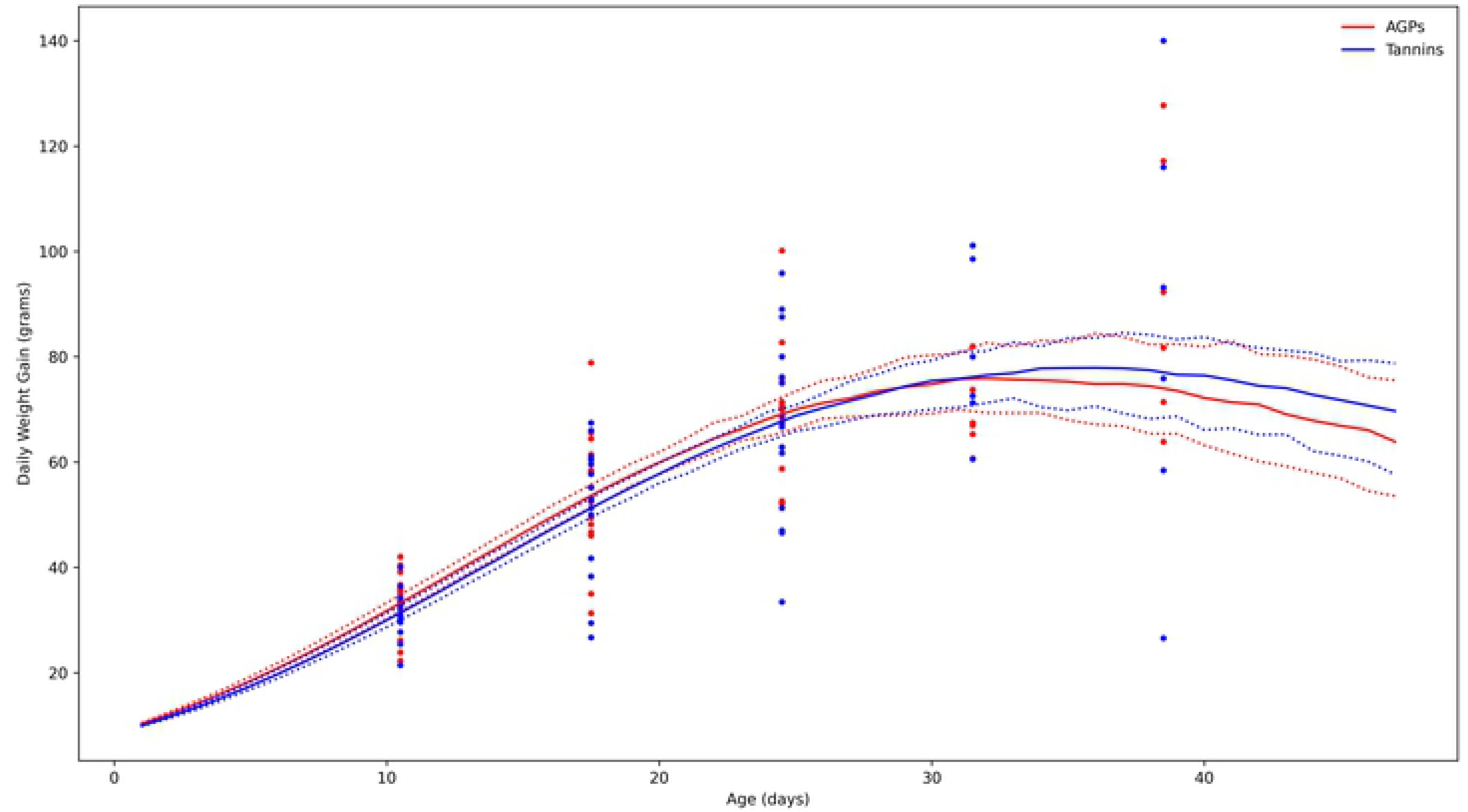
Daily weight gain derived from Gompertz equation. Solid lines represent estimated mean daily weight gains at different ages (R*Wf) for tannins (blue) and AGPs (red) treated flocks, dotted lines represents CI at 95% for the growth curve after 100000 montecarlo simulations. Circles represent average daily gains estimated from observed weekly weights (see table 4).

**Table 4:** Growth performance in broilers under AGPs or tannins programs (experiment 2).

| Age | AGPs | Tannins | p-values |
| --- | --- | --- | --- |
|  | Body weight (grs.)<sup>1</sup> |  |  |
| 7 days | 159.18 | 153.41 | 0.948 |
| 14 days | 402.35 | 383.57 | 0.959 |
| 21 days | 785.34 | 750.79 | 0.947 |
| 28 days | 1271.88 | 1229.68 | 0.858 |
| 35 days | 1801.96 | 1766.09 | 0.680 |
| 42 days | 2315.62 | 2305.73 | 0.514 |

<sup>1</sup> Values are expressed as mean.

Tannins/AGP free treated flocks showed a mild reduction (no significant) in total mortality (4.169% vs. 3.823%) and the mortality to the first week (0.873% vs. 0.721 %). During the second productive cycle a respiratory disease outbreak occurred in a barn under tannin treatment, increasing global tannins mortality (11.3%) compared to AGP (8.3%) treated flocks. Since these flocks were treated with antimicrobials, data from this period were omitted from analysis. Productive historical records and results from the studied period are summarized in figure 2.

**Figure 2:**
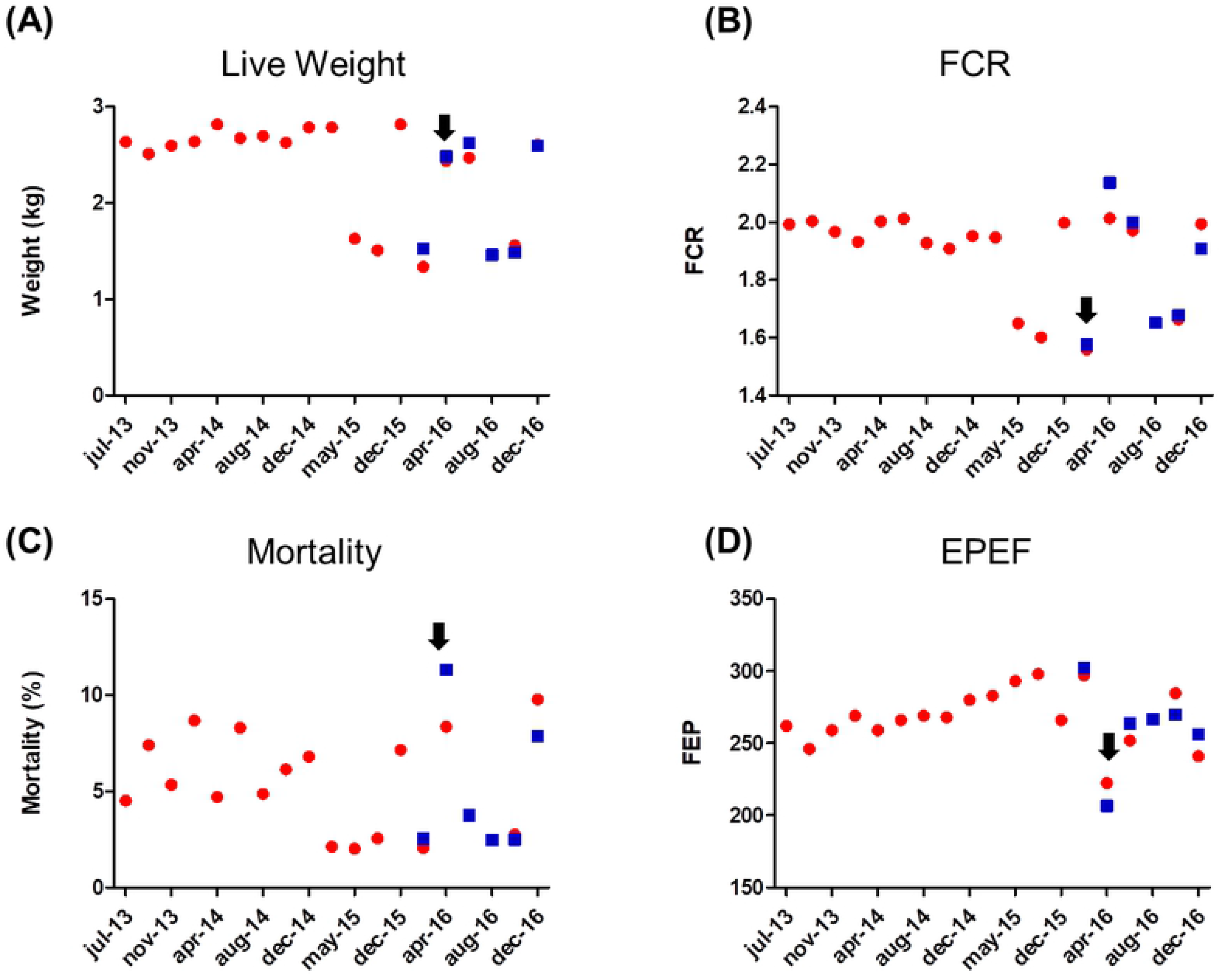
Productive historical records and results from experiment 2. A) Live body weight at the end of the productive cycle; B) Feed conversion ratio (FCR); C) Mortality; and D) European Poultry Efficiency Factor (EPEF). The period included in the present study starts from December 2015 (dec-15) to December 2016 (dec-16); red dots represent data from flocks under AGPs program and include data from previous productive cycles; blue squares represent data from AGPs free/tannins based program. Black arrow indicates the occurrence of a respiratory disease outbreak.

#### Necropsy findings

A total of 180 birds (with an average of 10 birds per flock/time) were evaluated. A description of intestinal lesions from this experiment is shown in table 5. In general, a mild reduction of gross intestinal lesions was observed among the tannin treated flocks examined on starter (day 21) and finisher periods (days 35 or 42). In contrast, lesion scores were higher in jejunum of birds from the tannin treated flocks during finisher periods. The condition of footpads deteriorated towards slaughter age, and despite this tendency was similar in both treatments, the tannin treated flocks (AGP free), showed reduced frequency (77.8% vs 64.4 %; p<0.05) of footpad lesions and reduced severity as represented by the mean score lesion (1.067 vs 0.766; p<0.0001).

**Table 5:** Frequency and score of intestinal gross lesions^1^ (Experiment 2)

|  | AGPs |  | Tannins |  |
| --- | --- | --- | --- | --- |
|  | Frequency (%) | Score (mean) | Frequency (%) | Score (mean) |
| **Day 21** |  |  |  |  |
| Duodenum | 2 | 0.28 | 2 | 0.24 |
| Jejunum | 16 | 0.74 | 4 * | 0.44 ** |
| Ileum | 4 | 0.62 | 6 | 0.62 |
| **Day 35-42** |  |  |  |  |
| Duodenum | 12.5 | 0.6 | 7.5 | 0.32 *** |
| Jejunum | 22.5 | 0.85 | 22.5 | 1.12 * |
| Ileum | 12.5 | 0.65 | ND | ND |
| **Total** |  |  |  |  |
| Duodenum | 6.7 | 0.42 | 4.5 | 0.27 |
| Jejunum | 18.9 | 0.78 | 12.2 | 0.74 |
| Ileum | 7.8 | 0.63 | 3.4 | 0.64 |

\*: $p < 0.05$ ; \*\*: $p < 0.01$ ; \*\*\*: $p < 0.001$ . ND: no lesions detected.
<sup>1</sup> Intestinal gross lesion score: 0: no apparent gross lesions; 1: removable fibrin deposit; 2: isolated focal necrosis or ulceration (1 to 5 foci); 3: multiple focal necrosis or ulceration (6 or more foci); 4: extensive areas of necrosis; 5: diffuse necrosis, presence of attached pseudomembrane)

#### Histomorphology

At day 21, reduced villus height in duodenum (p< 0.05) and jejunum (p< 0.001) were observed in tannin treated flocks, while ileal villi were higher (p< 0.001). In contrast, on finisher period villus height was increased in duodenum (p< 0.001) and jejunum (p<0.001). The means of duodenal, jejunal and ileal villus height, crypt depth, and villus:crypt ratio are presented in Table 6.

**Table 6:** Intestinal histomorphometry (Experiment 2)

|  |  | **AGPs** | **Tannins** | **p-values** |
| --- | --- | --- | --- | --- |
| **Day 21** |  |  |  |  |
| Duodenum | Villus height (µm) | 1602 ± 35 | 1488 ± 34 | 0.027 |
|  | Crypt depth (µm) | 187 ± 7 | 184 ± 9 | 0.812 |
|  | Villus/Crypt ratio | 9 ± 0.3 | 8.9 ± 0.5 | 0.906 |
| Jejunum | Villus height (µm) | 852 ± 16 | 765 ± 18 | 0.0007 |
|  | Crypt depth (µm) | 152 ± 5 | 149 ± 6 | 0.709 |
|  | Villus/Crypt ratio | 5.87 ± 0.2 | 5.64 ± 0.3 | 0.565 |
| Ileum | Villus height (µm) | 401 ± 15 | 501 ± 17 | <0.0001 |
|  | Crypt depth (µm) | 121 ± 3 | 138 ± 4 | 0.0028 |
|  | Villus/Crypt ratio | 3.35 ± 0.1 | 3.78 ± 0.2 | 0.068 |
| **Day 35 – 42** |  |  |  |  |
| Duodenum | Villus height (µm) | 1654 ± 23 | 1864 ± 31 | <0.0001 |
|  | Crypt depth (µm) | 144 ± 6 | 135 ± 6 | 0.312 |
|  | Villus/Crypt ratio | 12.64 ± 0.5 | 15.87 ± 0.9 | 0.016 |
| Jejunum | Villus height (µm) | 946 ± 22 | 1102 ± 28 | <0.0001 |
|  | Crypt depth (µm) | 158 ± 8 | 171 ± 5 | 0.211 |
|  | Villus/Crypt ratio | 7.27 ± 0.3 | 6.81 ± 0.2 | 0.295 |
| Ileum | Villus height (µm) | 516 ± 14 | 480 ± 12 | 0.065 |
|  | Crypt depth (µm) | 119 ± 4 | 118 ± 3 | 0.790 |
|  | Villus/Crypt ratio | 4.5 ± 0.1 | 4.2 ± 0.1 | 0.165 |

#### Cecal microbiota

Alpha diversity parameters did not show significant differences between the two treatments. Except for the Simpson’s index, diversity parameters showed a strong negative correlation with productive efficiency and these correlations were similar among treatments (Table S2). At the phylum level, the microbiota profiles also showed great similarity, with a marked predominance of species belonging to phylum Firmicutes. Relative abundance of the phyla Bacteroidetes and Tenericutes showed a negative correlation with productive efficiency (r: -0.52, p<0.05 and r: - 0.57, p<0.05 respectively) while a positive correlation was observed with Firmicutes (r: 0.48, p<0.05), these correlations were stronger within the conventional (AGP treated) groups. No significant differences were observed in the Firmicutes/Bacteroidetes ratio, although it was slightly higher in birds treated with tannins (Table S2). Significant differences were found in the relative abundance of ten bacterial taxa between AGP and tannin treated flocks (p<0.05), while four other taxa showed a trend (p<0.10) (Table 7a). Bacterial groups overrepresented in birds treated with AGP include the unclassified members of families *Desulfovibrionaceae* and *Synergistaceae*, genera *Slackia*, *Anaerostipes* and *Clostridium*, and species *Bacteroides coprophilus*, *Megamonas hypermegale* and *Escherichia coli*. On the other hand, in the birds treated with tannins, members of the family *Bifidobacteriaceae*, genera *Butyricimonas, Coprococcus*, *Ruminococcus* and species *Lactobacillus helveticus* and *Subdoligranulum variabile* were increased. Among these taxonomic groups, the genera *Butyricimonas* shows a positive correlation with BW and FCR, while the genera *Clostridium* was correlated with decreases in productive efficiency (Table 7b).

**Table 7a:** Cecal microbiota composition, AGPs vs tannins (Experiment 2)

| Family | Genus | Species | AGPs |  | Tannins |  | p-values |
| --- | --- | --- | --- | --- | --- | --- | --- |
|  |  |  | Rel. Ab.<br>(%) | Std. Dev.<br>(%) | Rel. Ab.<br>(%) | Std. Dev.<br>(%) |  |
| *Bifidobacteriaceae* | Unclassified |  | 0,20770 | 0,18275 | 0,77609 | 0,60826 | 0,009 |
| *Coriobacteriaceae* | *Slackia* | Unclassified | 0,06236 | 0,15305 | ND | ND | 0,001 |
| *Bacteroidaceae* | *Bacteroides* | *coprophilus* | 0,30551 | 0,66617 | ND | ND | 0,023 |
| *[Odoribacteraceae]* | *Butyricimonas* | Unclassified | 0,35844 | 0,35595 | 0,79575 | 0,61195 | 0,089 |
| *Lactobacillaceae* | *Lactobacillus* | *helveticus* | 1,19341 | 1,09405 | 2,93944 | 1,84471 | 0,020 |
| *Clostridiaceae* | *Clostridium* | Unclassified | 0,02676 | 0,04389 | ND | ND | 0,001 |
| *Lachnospiraceae* | *Anaerostipes* | Unclassified | 0,01892 | 0,03556 | ND | ND | 0,001 |
|  | *Coprococcus* | Unclassified | 0,16601 | 0,25298 | 0,54717 | 0,33977 | 0,013 |
|  | *[Ruminococcus]* | Unclassified | 0,39645 | 0,38623 | 1,00365 | 1,01141 | 0,083 |
| *Ruminococcaceae* | *Subdoligranulum* | *variabile* | 0,65751 | 0,57424 | 1,23472 | 0,78919 | 0,079 |
| *Veillonellaceae* | *Megamonas* | *hypermegale* | 0,31234 | 0,47607 | 0,02222 | 0,07025 | 0,100 |
| *Desulfovibrionaceae* | Unclassified |  | 1,03726 | 0,62765 | 0,24365 | 0,29632 | 0,006 |
| *Enterobacteriaceae* | *Escherichia* | *coli* | 0,13751 | 0,17889 | 0,01695 | 0,04001 | 0,074 |
| *Synergistaceae* | Unclassified |  | 0,10158 | 0,17299 | ND | ND | 0,006 |

**Table 7b:** Cecal microbiota composition correlated with productive indicators (Experiment 2)

| Family | Genus | Species | BW |  | FCR |  | EPEF |  |
| --- | --- | --- | --- | --- | --- | --- | --- | --- |
|  |  |  | R | p-values | r | p-values | r | p-values |
| *Bifidobacteriaceae* | Unclassified |  | 0.012 | n.s. | -0.084 | n.s. | 0.123 | n.s. |
| *Coriobacteriaceae* | *Slackia* | Unclassified | 0.139 | n.s. | 0.183 | n.s. | -0.358 | n.s. |
| *Bacteroidaceae* | *Bacteroides* | *coprophilus* | 0.213 | n.s. | 0.232 | n.s. | -0.338 | n.s. |
| *[Odoribacteraceae]* | *Butyricimonas* | Unclassified | 0.583 | 0.011 | 0.585 | 0.006 | -0.333 | n.s. |
| *Lactobacillaceae* | *Lactobacillus* | *Helveticus* | 0.069 | n.s. | 0.285 | n.s. | -0.260 | n.s. |
| *Clostridiaceae* | *Clostridium* | Unclassified | -0.014 | n.s. | 0.216 | n.s. | -0.554 | 0.011 |
| *Lachnospiraceae* | *Anaerostipes* | Unclassified | 0.19 | n.s. | 0.185 | n.s. | -0.199 | n.s. |
|  | *Coproccoccus* | Unclassified | 0.155 | n.s. | 0.306 | n.s. | -0.235 | n.s. |
|  | *[Ruminococcus]* | Unclassified | 0.17 | n.s. | 0.158 | n.s. | 0.062 | n.s. |
| *Ruminococcaceae* | *Subdoligranulum* | *variabile* | -0.44 | n.s. | -0.466 | n.s. | 0.434 | n.s. |
| *Veillonellaceae* | *Megamonas* | *hypermegale* | 0.257 | n.s. | 0.247 | n.s. | -0.172 | n.s. |
| *Desulfovibrionaceae* | Unclassified | Unclassified | -0.105 | n.s. | -0.168 | n.s. | 0.157 | n.s. |
| *Enterobacteriaceae* | *Escherichia* | *Coli* | -0.201 | n.s. | -0.244 | n.s. | 0.365 | n.s. |
| *Synergistaceae* | Unclassified | Unclassified | 0.209 | n.s. | 0.195 | n.s. | -0.180 | n.s. |

BW: body weight; FCR: Feed conversion rate; EPEF: European productive efficiency factor; n.s.: no significant

### Experiment 3

#### Necropsy findings

The groups treated with the mix of tannins; showed a clear reduction in the frequency and score of intestinal lesions observed on day 19, after an experimental challenge with a NE *C. perfringens* strain. Birds treated with tannins mix presented lower percentages of gross lesions in jejunum (43.75%, p<0.01) and ileum (18.75%; p<0.05), compared with the results obtained from positive control, 74.19% and 45.16% for jejunum and ileum respectively. The addition of individual tannins (chestnut or quebracho) had no effect on the frequency of gross lesions, although a no-significant reduction was observed in the incidence of severe lesions (score >3, data not shown). Reduced lesion score was observed in the jejunum (1.19 vs 1.97; p<0.01) and ileum of mix treated birds (0.63; p<0.05). Chestnut treated groups also showed a reduction in ileum gross lesions (0.58; p<0.05), in comparison with the score registered for the positive control group (1.29). Results from this trial are summarized in table 8. No intestinal gross lesions were detected in the negative control group (no challenge-no additives). Microscopic observation of these lesions revealed foci of necrosis, haemorrhage and epithelial desquamation. In most severe cases, accumulation of fibrinous exudates was observed. Mucosal smears of gross lesions in the small intestine of challenged groups showed abundant short Gram-positive bacilli compatible with *C. perfringens*. Bacteriological and molecular probes confirmed the identity of the obtained isolates (data not shown).

**Table 8:** Frequency and score of intestinal gross lesions^1^ (Experiment 3)

| Treatment | Jejunum |  | Ileum |  |
| --- | --- | --- | --- | --- |
|  | Frequency (%) | Score (mean) | Frequency (%) | Score (mean) |
| Positive control | 74.19 | 1.97 | 45.16 | 1.29 |
| Chestnut | 66.67 | 1.71 | 16.67* | 0.58* |
| Quebracho | 76.67 | 1.67 | 43.33 | 1.00 |
| Tannins mix | 43.75** | 1.19** | 18.75* | 0.63* |

\*: $p < 0.05$ ; \*\*: $p < 0.01$ .
<sup>1</sup> Intestinal gross lesion score: 0: no apparent gross lesions; 1: removable fibrin deposit; 2: isolated focal necrosis or ulceration (1 to 5 foci); 3: multiple focal necrosis or ulceration (6 or more foci); 4: extensive areas of necrosis; 5: diffuse necrosis, presence of attached pseudomembrane).

## DISCUSSION

The growing global concern about the emergence and spread of antimicrobial resistant microorganisms encouraged the search for alternatives that do not negatively impact animal health or production efficiency [1–4, 26]. Among described alternatives, phytochemicals and tannins in particular, are probably some of the most promising candidates since many studies assessed animal responses to the dietary inclusion of these compounds [7, 27, 28] and some of the reported biological activities are similar to those proposed for AGPs [5, 6]. Although the beneficial effects of phytochemicals as feed additives in poultry have been previously described [8, 29–31], and different tannin-based products have been used in commercial operations for many years now [5, 14], descriptions on their application as main component of an AGP free program under commercial conditions are scarce [6]. In the present study, results from two different field trials show that dietary addition of a blend of chestnut and quebracho tannins at 0.1% (w/w) represents a promising alternative for AGPs, since broiler chickens raised under commercial conditions achieve similar productive results either in AGP or tannins-based programs. In addition, necropsy findings and results from histology or microbiota studies described in this study provide evidence that this blend of tannins contributes to improve gut health status and reduce intestinal alterations associated with *C. perfringens* NE.

In the past, tannins have been considered harmful to monogastric animals due to their negative effect on nutrient digestibility, nitrogen retention and productive qualities [32, 33], but current research has led to an understanding of the chemical and biological activities of these complex compounds, transforming the conventional anti-nutritional concept of tannins [6-9, 29-31]. In fact, their effects have been shown to differ among animal species and to depend on time and dose exposure as well as on variations in chemical structures [28]. Previous works using tannins to promote growth show variable results, while the works performed by Manelli et al. [29] and Liu et al [30] describe positive effects on poultry growth after the addition of chestnut tannins (1-3 g/kg), Jamroz et al [34] did not find any significant effects on body weight gain and feed conversion in broilers supplemented with chestnut tannins (250 or 500 mg/kg), similar results are described by Brenes et al [35] in broilers fed with grape pomace (condensed tannins about 1,500 mg/kg) at different levels (15 mg, 30 mg, and 60 mg/kg). These differences may be related to the tannin source, manufacture conditions and other conditions not described in the mentioned studies. Although mechanisms involved in tannin induced growth promotion are not fully understood, beneficial effects of tannins can be related to diverse biological activities such as antioxidative properties, stabilization of intestinal microbiome and improvement of immune system through immunomodulatory effects [36–38]. In the present work, results from the histological evaluation show a differential response of the intestinal epithelium among AGPs and tannins treated birds, which may indirectly reflect some of the mentioned biological effects for tannins. However, the effect of tannins on enterocytes may be different depending on bird age and intestinal segment. Jamroz et al, [34] using similar dietary doses of tannins, describe disturbances in intestinal wall morphology and function suggesting potential cytotoxic effects, while other works in which lower doses of tannins are used describe proliferative and protective effects on enterocytes [39]. These apparent opposite actions indicate that potential benefits of tannins and other phytochemicals will depend on the effective concentrations of bioactive molecules that reach the epithelia and the presence or absence of metabolites produced by the resident microbiota. The bioavailability of tannins is an essential characteristic for their functionality in the gastrointestinal tract. However, bioavailability varies between different tannins depending on several factors, including molecular structure, molecular weight, composition of the derivatives, affinity to protein, etc. It is believed that absorption of these components is inversely correlated with their polymerization degree [40, 41]. Tannins with a low bioavailability would hypothetically show important antimicrobial activities, whereas highly bioavailable tannins would be more beneficial as antioxidant and anti-inflammatory agents.

All tannins share some common features which enable classification of these compounds in two main groups: hydrolysable tannins (gallotannines, ellagitannines, gallic acid and ellagic derivatives) and condensed tannins (non-hydrolysable) called procyanidins containing condensed carbon chain typical for flavonoids [27, 28]. Chestnut tannins can be hydrolysed in the small intestine and be readily adsorbed by the mucosa, disappearing before reaching the distal gastrointestinal tract [42], affecting several cellular processes [39]. In contrast, quebracho tannins have lower bioavailability and *in vitro* antioxidant capacity than chestnut tannins [43], but these extracts contain mainly condensed tannins which are not easily hydrolysed, retaining most of their biological activities along the entire gastrointestinal tract. In the present work, the combination of hydrolysable (chestnut) and condensed (quebracho) tannins in feed would affect homeostasis of intestinal epithelial cells through a combination of direct and indirect effects, which finally contributes to avoid negative effects described elsewhere [34, 35].

Many reports describe the role of gastrointestinal microbiota in host health and productive efficiency [44, 45], also most of these works relate modifications of resident microbial communities by feed additives such as AGPs or phytochemicals with the observed improvements in performance. Although results from previous studies show that antimicrobials reduce microbial diversity and tannins, as well as other phytochemicals, can generate opposite effects [13], in the present study no differences were observed between birds under AGP or tannin-based programs. These findings agree with previous results described for bacitracin and other AGPs from our group and other ones under experimental conditions [46–48]. Besides the lack of changes in global microbiota structure, changes in the relative abundance of certain taxa were described. Although no correlations with productive efficiency were found in the present study, most of the bacterial groups increased in tannins flocks were previously associated with growth promoting effects such as *Lachnospiraceae* and *Ruminococcaceae* at the family level [44, 45], or *Lactobacillus* or *Bifidobacterium* at genus level [49].

Along with negative productive consequences, antimicrobial restrictions also have been associated with increased incidence of infectious diseases like *C. perfringens* NE and *Campylobacter* shedding [3, 17, 50], resulting in poor zootechnical performance and increased condemnations at slaughter [51, 52]. In the present study, clinical necrotic enteritis cases were not identified during farm visits nor informed by farmers, but sub-clinical necrotic enteritis was regularly observed during systematic necropsies of apparently clinically healthy chickens within each flock of AGP and tannin treated animals. The estimated prevalence of subclinical NE was reduced in the tannin treated groups (10-25%) compared with a 25-45% in AGP groups, concomitantly with an important reduction in the score of intestinal lesions in animals of 21 days of age and reduction of undigested feed in the distal part of the small intestine. In another work with similar experimental design, authors reported no clinical or subclinical NE in conventional AGP programs but a prevalence of 30-50% in AGP-free programs based on essential oils [17]. In contrast with our study where tannins additives were applied during the whole productive cycle, in the mentioned work essential oils were only included during NE outbreaks in order to reduce negative consequences. Authors described that this treatment contributes to reduce NE associated mortality but failed to control productive losses. Although several works described the *in vitro* inhibitory effects of these phytochemicals against *C. perfringens* [53], the variable results obtained evidence the fact that despite many phytochemicals such as essential oils or tannins share biological activities (i.e. antimicrobial activity) [5, 10, 13], other biological activities and their combinations are vital to achieve consistent and reproducible productive results comparable with those obtained with AGPs, especially during an outbreak of an infectious disease like NE. In the particular case of chestnut and quebracho tannins, besides the inhibitory effects against *C. perfringens* and its toxins previously described [10–11], these phytochemicals display other multiple specific activities which can be related to improvement of growth efficiency and gut health [5, 13, 38, 41]. In addition, biological activities described before such as astringency, antioxidant [34, 39], intestinal microbiota modulation [13] and immunomodulation [36, 37], allow animals undergoing an infectious challenge which produces damage on the intestinal mucosa like NE, to recover faster, limiting the negative impacts on productive parameters. Therefore, besides the fact that tannins have significant antimicrobial activities against bacterial pathogens as *Salmonella* [54] and *C. perfringens* [10, 11], fungi, virus, and protozoa [55, 56], other actions on the host physiology seem to be important to explain the *in vivo* results. The experimental challenge with *C. perfringens* in an *in vivo* model of NE corroborated that the blend of tannins added to feed can significantly reduce intestinal lesions. These results visibly show a synergistic effect of chestnut and quebracho tannins, as the blend of these tannins was more efficient to control the occurrence of NE intestinal lesions than individual tannins. Also, the differences in the severity and distribution of intestinal gross lesions observed between the commercial blend and individual components suggest that each of the tested tannins made differential contributions to the control of NE. Following commercial blend, chestnut was the second most effective additive to control NE consequences. These observations can be explained in part by a direct and strong bactericidal effect [10], combined with differential activities along the different segments of the gastrointestinal tract as described before [34, 39, 41, 43]. While chestnut tannins can be hydrolysed in the small intestine and be readily absorbed by the mucosa, disappearing before reaching the distal gastrointestinal tract [42] with a probable reduction in their antimicrobial properties, derived metabolites may contribute to disease control through other cellular processes [39, 43], increasing epithelial resistance to pathogen actions and favouring recover. In contrast, quebracho extracts contain mainly condensed tannins which are not easily hydrolysed, retaining their bacteriostatic activity against *C. perfringens* [10] along the entire gastrointestinal tract and probably in the faeces/litter.

Searching for alternatives to AGPs has become a priority for the animal productive sector since several countries started limiting antibiotic use in animal production some years ago [4-7, 53, 57, 58]. Results from this study suggest that the use of tannin-based feed additives appears as an attractive alternative to promote gut health and productive efficiency, even as a main component of an AGP-free program. Additionally, necropsy observations during field trials and results from experimental *C. perfringens* challenge suggest that the blend of chestnut and quebracho tannins can help prevent and control NE with a reduction in the negative impact/consequences for poultry industry. Our data support the concept of replacing AGPs by phytochemical feed additives in commercial broiler flocks, preserving health and performance results. Since these natural products are complex substances with many bioactive compounds that do not leave residues in the carcass or other by-products and also reduce the chances of selection of antimicrobial resistance [11], their use is possible during the entire productive cycle without need for rotation. Moreover, since the farms enrolled in the present study followed management and vaccination procedures commonly used within the industry and also feed formulation did not differ from current industry standards, the present results can be functional for most of the existing broiler chicken production systems.

## Acknowledgements

This research was supported by Instituto Nacional de Tecnología Agropecuaria and the Consejo Nacional de Investigaciones Científicas y Tecnológicas-Argentina. We thank, Ms. Laura Gonzales, Ms. Jesica S. Bucci, Mr. Ignacio de la Fuente, Mr. Facundo Balbiani, Dr Johana Dominguez, Dr. Natalia Casanova, Dr. Pablo Chacana and Dr. Fernando Delgado for invaluable help at the lab. We thank Mrs. Daniela Losada-Eaton for her assistance in reviewing the manuscript.

## Authors’ contributions

EAR, LMR, CC, VG, ML and MEFM participated in the design of the field trials, also perform necropsies and sample collection from the mentioned trials. EAR, LMR and MEFM design and perform experiments and necropsies associated with the *C. perfringens* NE model. EAR perform the histological analysis. JMDC perform the microbiota analysis. OAB, LMR and MEFM analyzed the data. EAR, LMR and MEFM wrote the paper. All authors contributed to the critical revision of the manuscript for important intellectual content and have seen and approved the final draft. All authors read and approved the final manuscript.

